# A 3D human liver model of nonalcoholic steatohepatitis

**DOI:** 10.1101/2020.02.07.938787

**Authors:** Marion Duriez, Agnes Jacquet, Lucile Hoet, Sandrine Roche, Marie-Dominique Bock, Corinne Rocher, Gilles Haussy, Xavier Vigé, Zsolt Bocskei, Tamara Slavnic, Valérie Martin, Jean-Claude Guillemot, Michel Didier, Aimo Kannt, Cécile Orsini, Vincent Mikol, Anne-Céline Le Fèvre

## Abstract

We have developed an *in vitro* preclinical 3D Non-Alcoholic SteatoHepatitis (NASH) model by co-culturing four human primary liver cell types: hepatocytes, stellate, endothelial and Kupffer cells. Cells were embedded in a hydrogel of rat collagen in 96-well plate and a NASH-like environment was induced with a medium containing free fatty acids (FFAs) and tumor necrosis factor α (TNFα). This model was characterized by biochemical, imaging and transcriptomics analysis. On the one hand, we succeed in defining suitable culture conditions to maintain the 3D co-culture up to 10 days *in vitro* with the lowest level of steatosis, and reproducible low levels of inflammation and fibrosis. On the other hand, we induced NASH disease with a custom medium mimicking NASH features (hepatocyte injury, steatosis, inflammation and fibrosis). The 10-day cell viability and cost effectiveness of the model make it suitable for medium throughput drug screening and provide attractive avenues to better understand disease physiology and to identify and characterize new drug targets.

**Summary:** We developed a 3D human liver model which exhibits many features of non-alcoholic steatohepatitis and that could become a platform for medium throughput drug screening.

## Introduction

Non-alcoholic fatty liver disease (NAFLD) is an umbrella term that comprises a large spectrum of liver injuries varying in severity and leading to fibrosis. Among these, non-alcoholic fatty liver (NAFL) refers to hepatic steatosis alone, which is very common (Younossi et al., 2019) and driven by the accumulation of intracellular lipid droplets. Furthermore, non-alcoholic steatohepatitis (NASH) is defined as a more serious pathogenesis with inflammatory foci, hepatocyte damage, and fibrosis. Adverse hepatic outcomes related to NASH may include cirrhosis, liver failure and hepatocellular carcinoma (Friedman et al., 2018). NAFLD is associated with obesity and features of metabolic syndrome, including hypertension, dyslipidemia, central adiposity, insulin resistance or diabetes (Buzzetti et al., 2016, Younossi et al., 2016). NASH with advanced fibrosis has been linked to increased overall and liver-related mortality (Dulai et al., 2017).

Today bariatric surgery is the most efficient procedure to reverse NASH and fibrosis in obese patients (Lassailly et al., 2015). Also, weight loss induced by diet and exercise has been shown to be effective in resolving NASH and improving hepatic fibrosis (Vilar-Gomez et al., 2015). Despite a marked increase in prevalence, NASH is still an orphan disease with no approved drugs. Thus, a better management of NAFLD and the development of new drugs and therapeutic options are urgently needed (Rinella and Sanyal, 2016). Although there has been steady progress in understanding NASH pathogenesis, the identification of therapeutic targets and the advancement of drug development have shown limited progress, mainly due to the lack of predictive preclinical models. Several animal models have been developed to study NAFL and NASH, but they do not accurately depict the human pathology, presumably because of NAFL/NASH heterogeneity (Santhekadur et al., 2018).

In drug discovery, hepatic *in vitro* models have been used to assess drug clearance and hepatotoxicity by investigating metabolism, enzyme induction and transporter function. Monolayer cultures of isolated primary rat or human hepatocytes remain the main investigative tools for drug testing. These 2D models have shown several limitations including a short lifetime and loss of function, likely resulting from dedifferentiation of primary hepatocytes (Godoy et al., 2013). Precision-cut liver slices, which contain primary hepatocytes, but also liver non parenchymal cells (NPCs), have also shown reduced lifetime, thus impairing the development of liver chronic disease models and robust drug testing.

More recently, 3D cultures have been developed to improve cell survival and to provide a more natural tissue-like environment. In the liver, the extracellular matrix (ECM) behaves as a scaffold for surrounding cells and *in vitro* artificial matrix aims to support ECM functions by promoting cell adhesion, cell differentiation and cell-to-cell communication (Martinez-Hernandez and Amenta, 1993, Wells, 2008, Abedin and King, 2010). 3D cultures of hepatic cell lines or human primary hepatocytes embedded in artificial scaffolds were shown to modify gene and cell surface receptor expression toward more mature-like phenotypes, resulting in the maintenance of hepatocyte polarization and functionality. Furthermore, it has been shown that collagen gels enhanced mechanical properties with good cell adhesion and a high survival rate for hepatocytes (Godoy et al., 2013). Thus, 3D cell culture systems appear more representative of hepatocyte physiology in liver tissue and provide opportunities to develop extended *in vitro* models of NASH and NAFLD.

The development of *in vitro* human 3D models to mimic liver architecture has been undertaken by several groups, especially for drug safety assessment (Oseini et al., 2018). It includes layered co-cultures, co-cultures on micro-patterned surfaces, spheroids and bioprinted liver tissue (Oseini et al., 2018). Many of these models are grown within specialized microfluidic devices to provide nutrients and oxygen transport (Oseini et al., 2018). Few 3D co-culture studies were designed as NAFL or NASH disease models to display key pathogenic phenotypes. Thus, steatosis was observed in NAFLD models supplemented with high concentrations of oleic and palmitic acids. Increased lipid accumulation was associated with altered gene expression and activity of several CYP450 enzymes. Only limited cytokine release was reported, likely due to the absence of Kuppfer cells in these *in vitro* systems. Therefore, such 3D models highlighted the need to co-culture additional cell types that could further incorporate features of inflammation and fibrosis and better reflect disease progression.

Feaver et al. set up a 3D model with primary hepatocytes, monocyte-derived macrophages and hepatic stellate cells (HSC) in hemodynamic and transport conditions. Correlations between the *in vitro* model and human biopsies were evidenced by transcriptomics, lipidomics and functional analysis (Feaver et al., 2016). This model requires differentiating monocytes into macrophages, which is time consuming, and yields only M2 macrophages, a non-physiological macrophage sub-population. In addition, this hemodynamic system is awkward to use for high throughput screening.

In this paper, we report on a new 3D co-culture model combining primary hepatocytes with hepatic stellate cells, sinusoidal endothelial cells and Kupffer cells. Activated hepatic stellate cells play a principal role in fibrosis initiation and development through the production of collagen, while Kupffer cells are involved in liver damage and inflammatory processes. Endothelial cells were more recently shown to play a pivotal role in NAFL/NASH progression (Ramachandran et al., 2019). This model was characterized biochemically, transcriptionally and displayed some key features of hepatic injury, steatosis, inflammation and early fibrosis. Its 10-day cell viability, as well as its reasonable cost effectiveness, make it compatible for medium throughput screening in 96 well plates. This co-culture model thus provides a valuable platform to better understand NASH disease progression, and to evaluate drug targets and compound activity.

## Results

### Human liver primary cells 3D co-culture characteristics in healthy conditions

Setting up a human *in vitro* NASH disease model requires developing a co-culture containing the different cells involved in the pathogenesis and maintaining it over an extended period of time in culture to induce this chronic disease. We developed a 3D co-culture with primary cells using the RAFT™ biological rat collagen hydrogel embedding human hepatocytes (PHH), hepatic stellate cells (HSC), Kupffer cells (KC) and liver endothelial cells (LEC). The addition of LEC as feeder cells in the 3D liver co-culture allowed us to stabilize the model by improving cell viability from one week up to two weeks (data not shown). PHH, HSC and LEC were seeded in the hydrogel of collagen at a 5:1:1 ratio respectively, reproducing the human liver ratio, and cultured in a 96 well plate in a homemade cell culture media named healthy media (**Fig. 1A)**. At Day 6, Kupffer cells were added to the co-culture since they did not tolerate the embedding process (data not shown) but were able to enter the hydrogel once formed. Several media were tested to maintain viability of the four cell types. The homemade Liver Co-culture optimized media, described in the Materials and Methods section, allowed us to obtain a viable co-culture of four cell types up to Day 10 (**Fig. 1B)**.

**Fig. 1:**
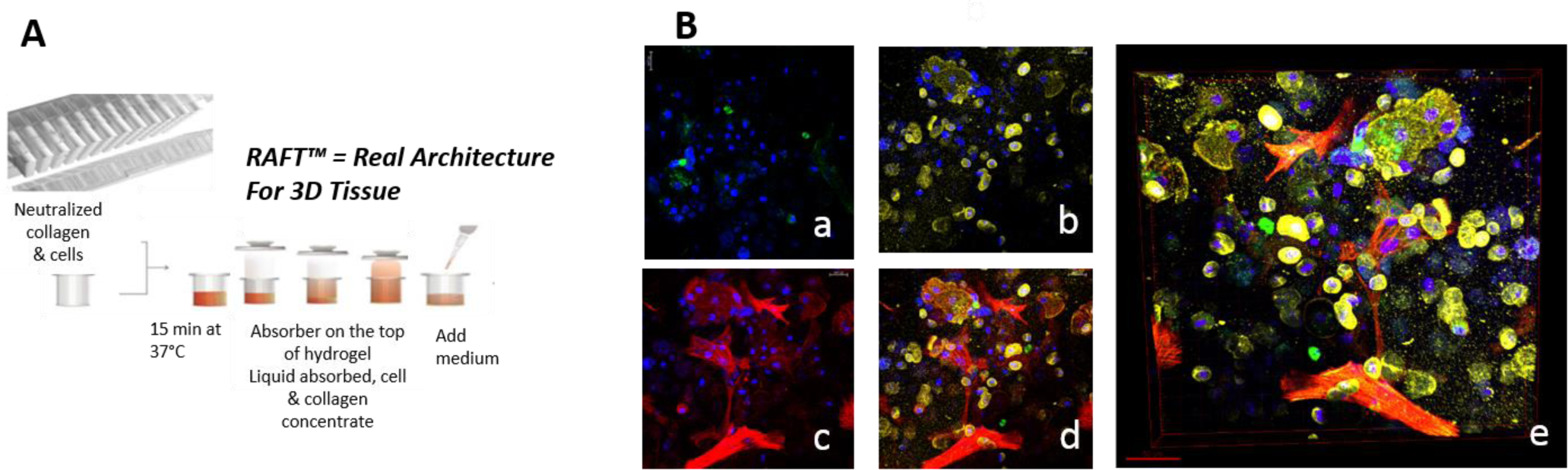
Set-up of human liver primary co-culture in a 3D environment. **(A)-** Real architecture for 3D model with RAFT™ system methodology illustration. **(B)-** PHH-KC-HSC-LEC co-culture at 10 days in 3D collagen matrix with healthy media. **(a)-** KC staining with CD68 in green and nucleus labeled in blue **(b)-** hepatocyte staining with CK18 in green and nucleus labeled in blue **(c)-** HSC staining with α-SMA in red and nucleus labeled in blue **(d)-**merge **(e)-** 3D reconstitution with IMARIS software 9.1.2

To assess the suitability of the model, co-culture viability, cytokine secretion and lipid droplet content were measured from Day 3 to Day 15 on four independent experiments. The human 3D liver co-culture at a basal level displays a good viability from Day 3 to Day 15 as reflected by ATP level measurement (**Fig. 2A**). PHH CYP3A4 enzymatic function was evaluated and it remains stable from Day 3 up to Day 13 (**Fig. 2B**). Inflammatory cytokines and chemokines IL-6, CXCL8 and CXCL10 were measured in co-culture supernatants. Basal levels of secreted IL-6 and CXCL8 remained stable from Day 3 to Day 15 with a non-significant increase at Day 8 resulting from the addition of Kupffer cells at Day 6 (**Fig. 2, C and D**). CXCL10 was almost undetectable before Day 8, after Kupffer cells addition, and its secretion in cell supernatant remained stable from Day 8 to Day 15 (**Fig. 2, C to E**). The triglyceride content of the human liver 3D co-culture was quantified, and PHH’s lipid droplets were observed by immunofluorescence by a green lipitox and a cytokeratin-18 co-staining (**Fig. 2, F and G, respectively**). A high basal level of triglyceride with no significant variation was detected from Day 3 to Day 15 (**Fig. 2F**). This result was correlated with the observation of an important intracellular lipid droplets staining in PHH from Day 3 to Day 10 (**Fig. 2G**). Of note is that the PHH triglyceride content in human liver 3D co-culture was higher than expected. Several media were tested on PHH, including William’s E media. All of them induced an elevated intracellular lipid droplet content (**Fig. S1**).

**Fig. 2:**
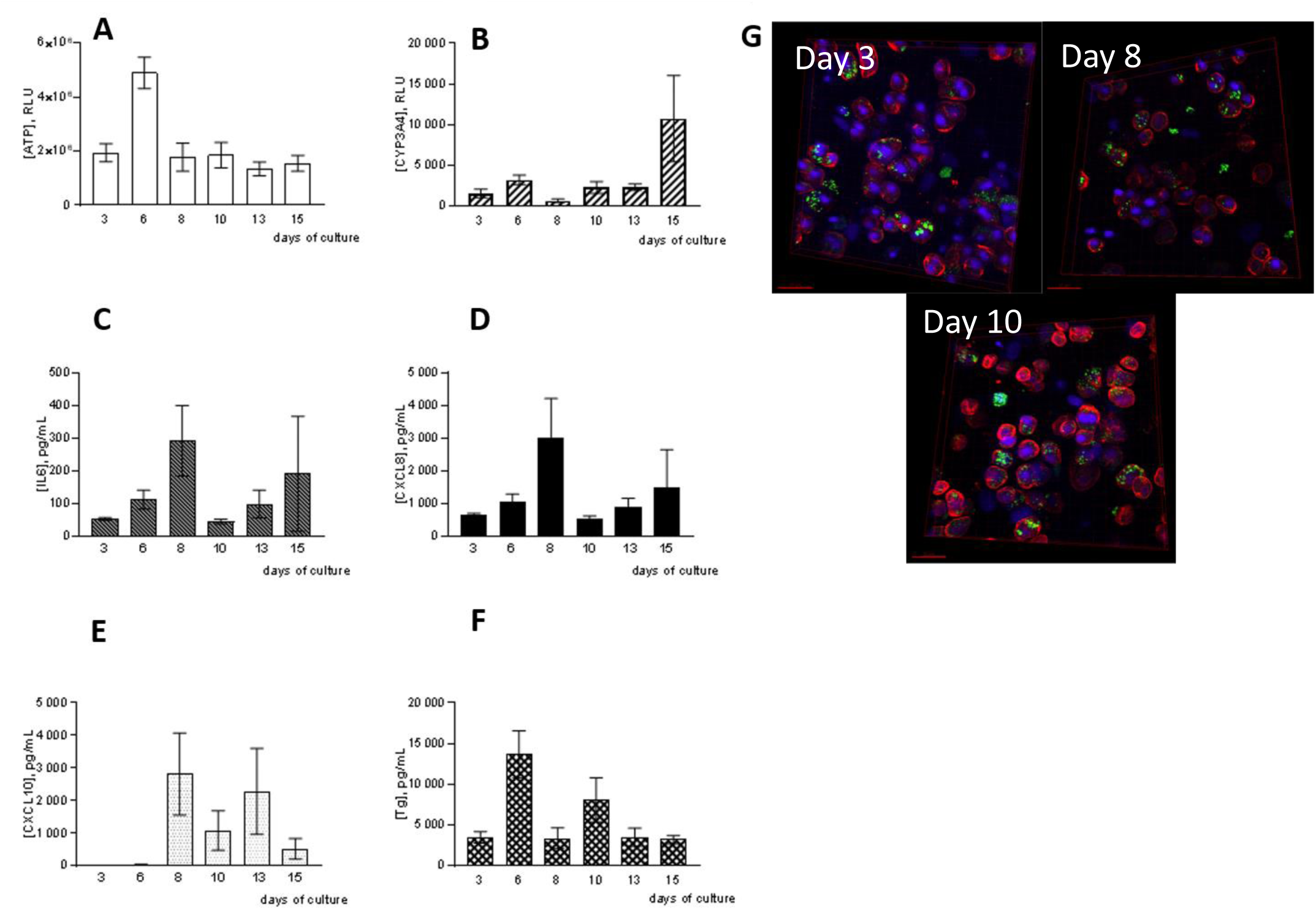
Human 3D liver primary co-culture characterization. **(A)** ATP intracellular in RLU. **(B)** CYP3A4 activity in RLU. (**C)** IL-6 (**D)** CXCL8 and **(E)** CXCL10 in pg/mL. (**F)** Tg content in pg/mL. (**G)** Kinetic of lipid droplet content in liver co-culture in 3D collagen matrix, healthy condition. PHH staining of CK18 in red, nuclear in blue and lipid droplet in green. *p< 0,050** p<0,010 ***p<0,001, two-way ANOVA with random effect followed by Bonferroni-Holm correction

Together, these results support a long-term viability and the stable expression of NASH-induced key features that can be directly quantified in the human 3D liver primary co-culture under healthy conditions up to 15 days (**Table S1**).

A transcriptomic analysis of human 3D liver primary co-culture under healthy condition was performed. Gene expression analysis was only performed on samples from day 3, 8 and 10 (n=3/condition) as the total RNA quantity extracted at day 13 and day 15 was too low for gene expression studies.

The expression of genes related to PHH activities, including the efflux transporters ABCC2, ABCBC1 and ABCB11, was first investigated. These genes are stably detected during the time course of the co-culture until Day 10 (**Table S2**) as well as UGT1A1 a gene encoding phase II UDP-glucuronosyltransferase enzyme, with a fold change <2 and/or with a p-value>0.05. Furthermore, CYP7A1 cytochrome P450 gene was expressed from Day 3 to 10 with no significant difference **(Table S2**), CYP3A4 expression, which is decreased at Day 8 and 10 compared to Day 3, is stable from Day 8 to 10 (**Table S2**) and was previously observed to be functional from Day 3 to 13 in 3D co-culture (**Fig. 2B).** Together, these genes reflect the polarization, functionality and metabolic activity of mature PHH in human 3D liver primary co-culture up to 10 days *in vitro*.

As expected, the expression of IL-6, CXCL8 and CXCL10 inflammatory cytokine genes is observed. IL-6 expression is stable between Days 3 and 10 (**Table S2**), whereas CXCL8 and CXCL10 are significantly down and up regulated from Days 3 to 8, resulting from the addition of Kupffer cells, but remain stable from Day 8 to 10 (**Table S2**). IL-6, CXCL8 and CXCL10 gene expression results are in line with their secretion observed in 3D co-culture supernatants (Fig. C, D and E). The expression of Col1A1 gene, encoding collagen compounds remains stable from Day 3 to Day 10 (**Table S2**), whereas ACTA2 gene, encoding α-SMA protein, slightly increases with a 2.8- and 2.4-fold change at Days 8 and 10, compared to Day 3, but remains stable from Days 8 to 10 (**Table S2**). ACTA2 gene increase from day 3 to 8, probably resulting from addition of KC since the latter were shown to activate HSC through soluble factors (Kolios et al., 2006). Together these results showed that HSC are not activated by a bystander effect of the co-culture under healthy conditions.

We succeeded in defining suitable culture conditions, including medium composition and chronological cell addition to maintain the four liver primary cell types in 3D co-culture up to 10 days *in vitro*. Furthermore, these culture conditions gave the lowest level of steatosis, together with reliable low levels of inflammation and fibrosis.

### Human 3D liver NASH model displays PHH injury and steatosis

The human *in vitro* 3D NASH model was set up by culturing the human primary 3D liver co-culture with a custom medium mimicking a NASH-like disease environment added from Day 3. This medium contains free oleic and palmitic fatty acids at 100 μM with a 2:3 ratio, as well as TNFα at 5 ng/ml.

The ATP levels measured in the human 3D liver co-culture did not show a significant difference between the healthy and NASH culture conditions and reflected a good viability which was maintained until Day 10 (**Fig. 3A**). ABCC2 and ABCB11 gene expression is not modulated during the co-culture, neither in NASH nor in healthy models, showing a stable PHH mature phenotype up to 10 days *in vitro* (**Fig. 3B** and **Table S3**). Cytochrome P450 CYP3A4, the expression of which is known to be reduced in NASH, displayed a reduced activity and gene expression in the NASH model as compared to healthy co-culture but without reaching statistical significance (**Fig. 3C**). However, at Day 10, three out the four experiments showed a decreased CYP3A4 activity (**Fig. 3C**), which correlated to a reduced gene expression at Day 10 (**Fig. 3D** and **Table S3**).

**Fig. 3.**
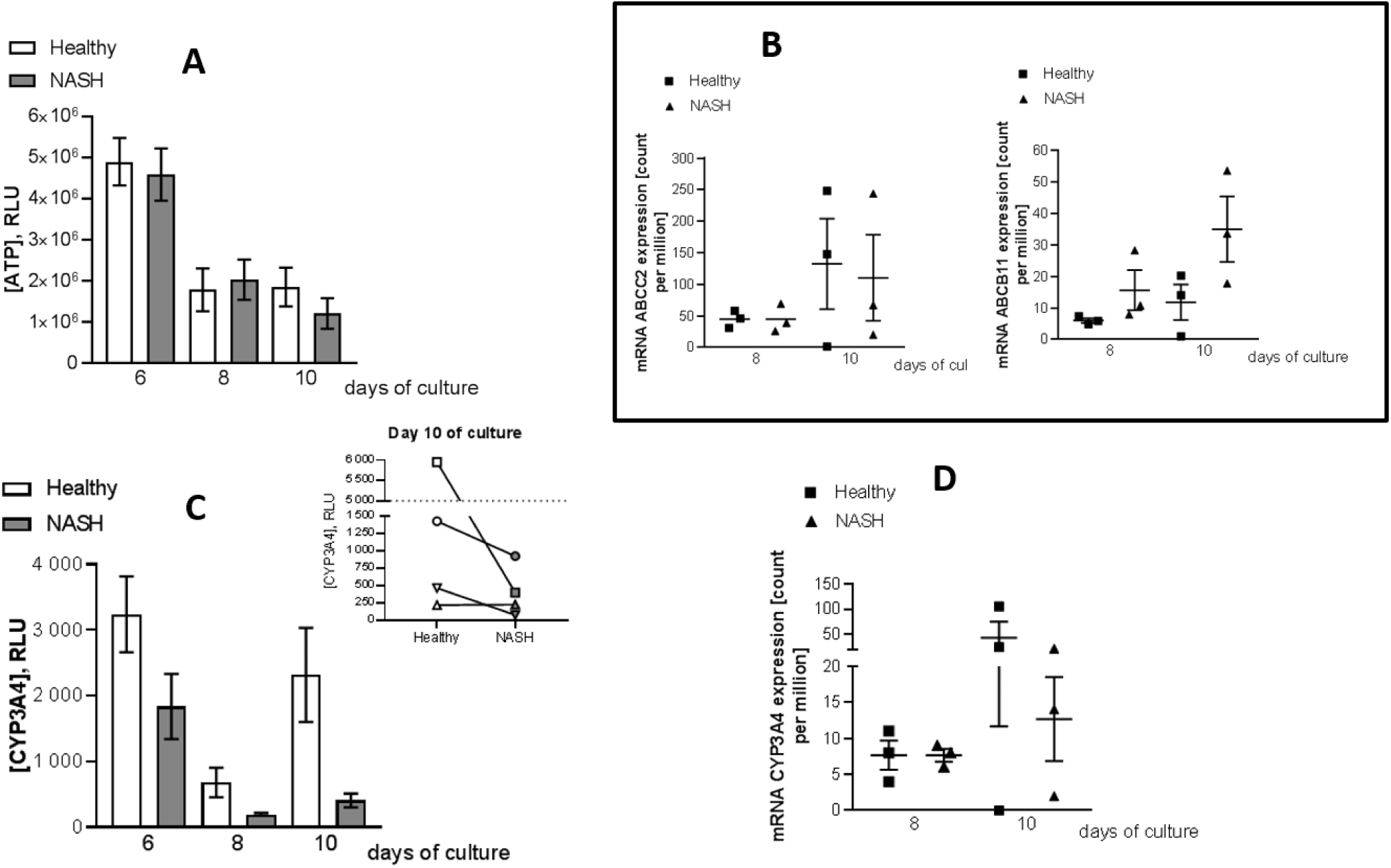
PHH injury characterization in human 3D liver NASH model. **(A)** Time course of ATP mean concentration (in RLU, +/-SEM) in 3D healthy and NASH models. (**B)** ABCC2 (left panel) and ABCB11 (right panel) mRNA expression in counts per million at Days 8 and 10 in healthy and NASH culture. (**C)** Time course of CYP3A4 mean concentration (in RLU, +/-SEM) in 3D healthy and NASH models (left panel) with a focus on the fourth independent experiment trend on Day 10 (right panel, each symbol represents an independent experiment). (**D)** CYP3A4 mRNA expression in counts per million at Days 8 and 10 in healthy and NASH culture conditions. *p< 0,050** p<0,010 ***p<0,001, Student’s test and DESeq2, respectively for panel A&C and panel B&D.

PHH steatosis assessed by triglyceride quantification in co-culture does not show significant differences between healthy and NASH culture conditions (**Fig. 4A**). Intracellular lipid droplets staining is observed in both conditions but without noticeable differences at Day 8 and 10 (**Fig. 4B**). As mentioned before, PHHs display a high basal level of triglycerides. This feature was observed for all the media tested, promoting PHH viability in the 3D collagen matrix and is aligned with biochemical quantification of steatosis.

**Fig. 4.**
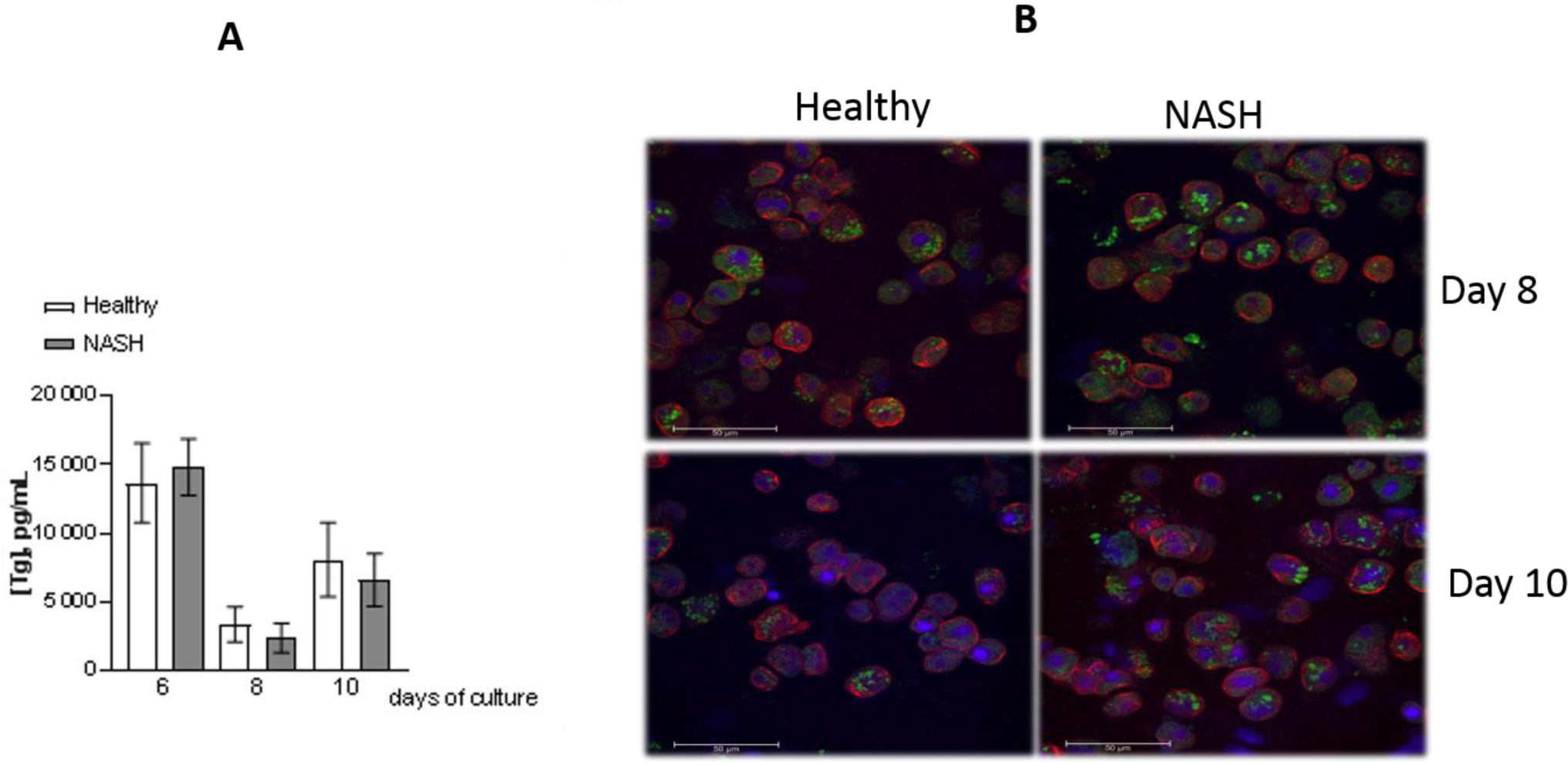
Assessment of steatosis feature. (**A)** Time course of triglyceride mean concentration content (in pg/mL, +/- SEM) in 3D healthy and NASH models. **(B)** Lipid droplets staining in healthy and NASH culture conditions (left and right panels respectively) at Day 8 and 10 (upper and lower panels respectively). Lipid droplets are stained in green, nucleus in blue and CK18 staining at PHH cell surface in red. Images acquired with a Leica SP8X confocal microscope. *p< 0,050** p<0,010 ***p<0,001, Student’s test.

The human *in vitro* 3D NASH model displays a similar stable viability as observed in the healthy 3D liver co-culture model, whereas it shows a decreased CYP3A4 activity in three out four experiments. Lipids droplets, characterizing steatosis, are observed in both conditions: the high basal level in healthy co-culture does not allow detection of an increase of triglyceride content in NASH conditions.

### Human 3D liver NASH model express inflammatory and tissue remodeling factors

The ability of human 3D liver co-culture to react to a pro-inflammatory environment was explored. The secretion of CXCL8, IL-6, CXCL10 and CCL2 in supernatants of healthy and NASH 3D co-culture were quantified by a multiplex assay at Days, 6, 8 and 10. TNFα (5 ng/mL) was added to the medium and changed every 2-3 days to induce an inflammatory process.

IL-6 secretion had a significant 5-fold increase at Day 6 in the 3D NASH model, compared to healthy condition (**Fig. 5A** and **Table S4**) and a significant increase of gene expression at Days 8 and 10 (**Fig.5B.**). The secretion of CXCL8 was significantly up-regulated in NASH co-culture at Day 6 and Day 8 with a 10.3- and 4.7-fold increase, respectively (**Fig. 5A** and **Table S4**). This observation correlated to an up-regulation in the expression of CXCL8 in 3D NASH co-culture from Day 8 to Day 10, with a respective fold increase of 13.4 and 14.6 (**Fig. 5B**). Finally, CXCL10 secretion is also significantly up-regulated in the 3D NASH model: 32.5 and 22.6-fold at Day 6 and Day 10, respectively (**Fig. 5A and Table S4**). A significant 8.7-fold increase of CCL2 at Day 6 is also observed in co-culture supernatant in the 3D NASH model (**Fig. 5C** and **Table S3**).

**Fig. 5.**
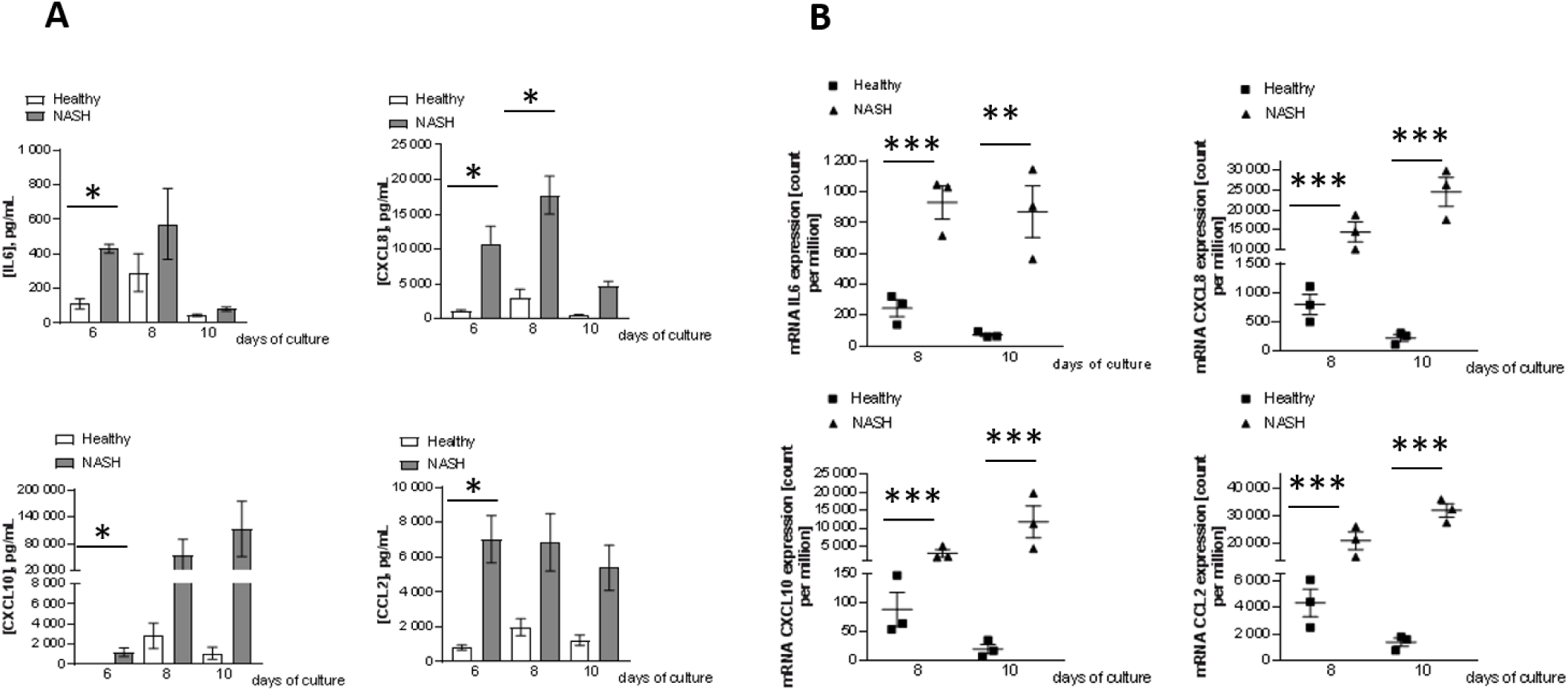
Inflammatory environment induced in 3D liver NASH model. **(A)** Time course of secreted IL-6, CXCL8, CXCL10 and CCL2 mean concentration (in pg/ml+/- SEM) at Days 6, 8 and 10 in healthy and NASH models. **(B)** IL-6, CXCL8, CXCL10 and CCL2 mRNA expression in count per millions at Days 8 and 10 in healthy and NASH conditions. *p< 0,050** p<0,010 ***p<0,001, Student’s test and DESeq2, respectively for panel A and panel B

The expression of the metalloproteinases MMP2 and MMP9 involved in tissue remodeling, and markers of early fibrotic events was also explored. MMP2 secretion in human 3D NASH co-culture significantly increased by 1.2, 1.7 and 1.4-fold at Days 6, 8 and 10 respectively (**Fig. 6A** and **Table S4**) and correlated with a significative upregulation in MMP2 gene expression at Day 10 (**Fig. 6B** and **Table S3**). Finally, MMP9 transcripts significantly increased both at Days 8 and 10, with a 3.2 and 6.6-fold change in the NASH environment (**Fig. 6C** and **Table S3**).

**Fig. 6.**
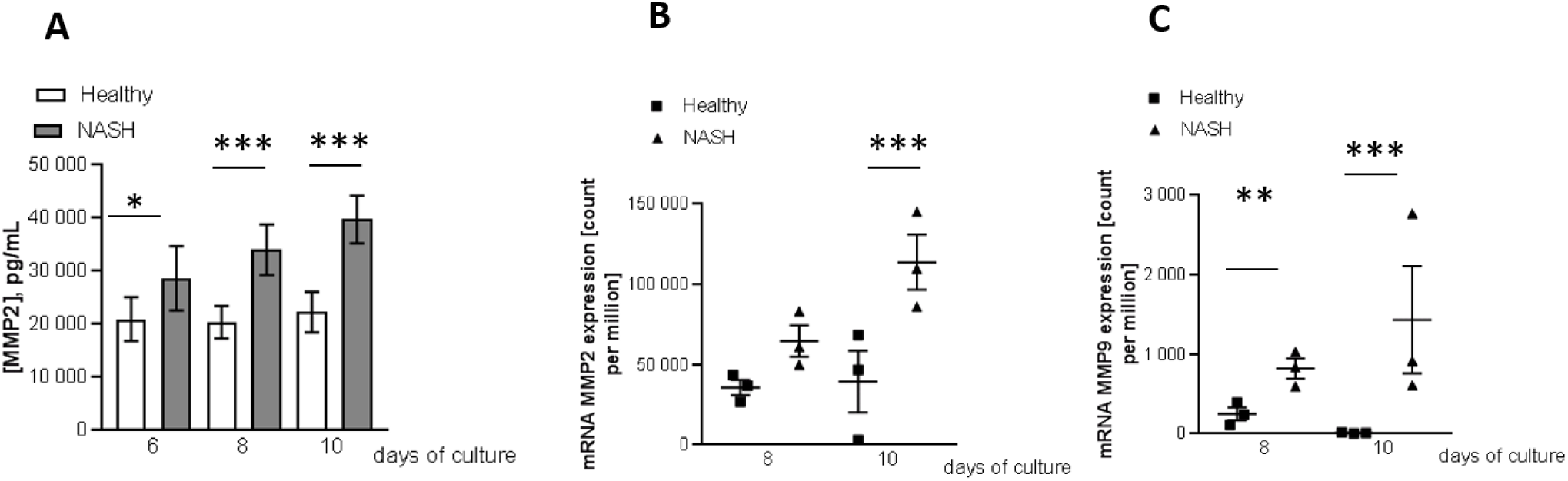
Early fibrotic tissue remodeling factors induced in 3D liver NASH model. **(A)** Time course of secreted MMP2 mean concentration (in pg/mL, +/- SEM) at Days 6, 8 and 10 in healthy and NASH models. **(B)** MMP2 and **(C)** MMP9 mRNA expression level in count per millions at Days 8 and 10 in healthy and NASH conditions. *p< 0,050** p<0,010 ***p<0,001, Student’s test and DESeq2, respectively for panel A and panel B&C.

Together these results show that human *in vitro* 3D NASH model is able to simulate a stable inflammatory and early fibrotic environment with the secretion of IL-6, CXCL8, CXCL10, CCL2 and MMP2 and MMP9 expression, respectively, as compared to healthy model.

### Human 3D NASH model regulates gene set related to pathways involved in NASH development

To better characterize the human *in vitro* 3D NASH model, a global gene expression analysis has been performed by RNAseq. For this purpose, three samples from day 3, 8 and 10 healthy 3D co-cultures and from day 8 and 10 *in vitro* 3D NASH model were processed.

Global RNA-seq datasets from 3D healthy and NASH models were first interrogated using a Principal Component Analysis (PCA) to cluster these two models during *in vitro* development (**Fig. 7A**). A distinct sample separation is observed between 3D healthy and NASH models (represented by circles and triangle, respectively), highlighting that NASH culture condition alters the overall gene expression pattern of 3D liver co-cultures (**Fig. 7A**). A time effect on both healthy and NASH 3D liver model is also visualized, being more noticeable between Days 8 and 10 in the 3D NASH co-culture.

**Fig. 7.**
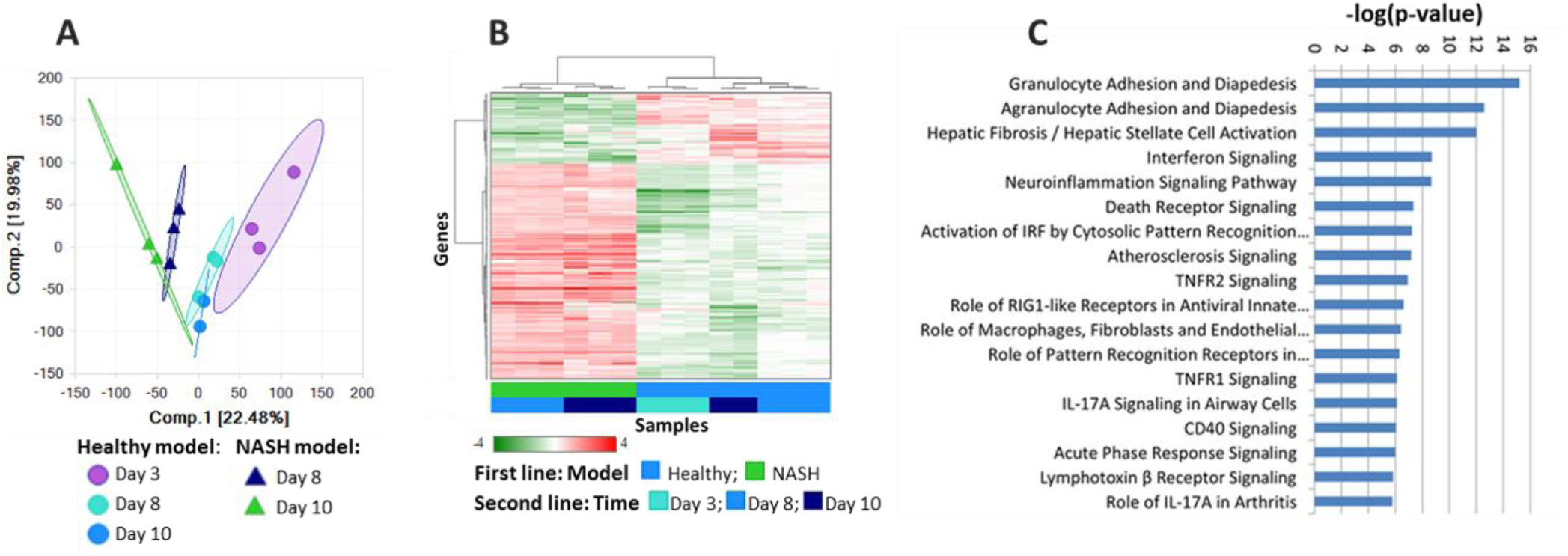
RNA-seq analysis of NASH and healthy 3D models. (A) Principal component analysis (PCA) for gene expression in the 3D model of NASH versus healthy state at Day 3, Day 8 and Day10. The PCA was performed using DEseq2 normalized expression data. (B) Clustering analysis of DGEs between the 3D model of NASH disease and biological healthy model, at Day 8. Heat map illustrating unsupervised hierarchical clustering of the 468 specifically regulated in NASH model vs biologically healthy model at Day 8 (log2 of DEseq normalized data). (C) Enrichment analysis of molecular pathways in NASH model at Day 8. Visualization of top 18 enriched canonical pathways in human NASH patients as compared to normal controls. Values are expressed as −log(p-value).

To identify pathways modulated under NASH-inducing conditions, differentially expressed genes (DEGs) between 3D healthy and NASH models were identified. A total of 659 significant DEGs were detected between 3D healthy and NASH at day 8 (**Fig. S2**). A gene subset was already modulated between Day 3 and Day 8 in the healthy 3D model and thus could not be assigned to NASH culture condition. In the end, we selected 468 DEGs in the 3D NASH model that were not found regulated over time in the healthy model. This list included 351 up-regulated and 117 down-regulated genes (**Fig. S2**). A hierarchical clustering analysis of healthy and NASH 3D liver models using these 468 DEGs confirmed a good separation of the samples from healthy and NASH 3D co-culture conditions (**Fig. 7B**). In addition, most of the DEGs after 8 days *in vitro* are still differentially regulated at Day 10.

We ran further pathway analysis of the 468 gene set differentially regulated in NASH conditions. Interestingly, among the top 18 enriched pathways, genes involved in hepatic fibrosis/hepatic stellate cell activation were found modulated, and this pathway was ranked in third position (**Fig. 7C**). The first two most strongly modulated gene clusters are associated with the activation of immune cell adhesion and the inflammatory process via the diapedesis pathways (**Fig. 7C**).

More precisely, genes encoding for claudin adhesion proteins, together with members of the immunoglobulin superfamily such as ICAM1, ICAM2 and VCAM1 were upregulated (**Fig. 8**). Other additional up-regulated inflammation/immune pathways included interferon signaling, IRF activation TNFR and IL17 signaling.

**Fig. 8.**
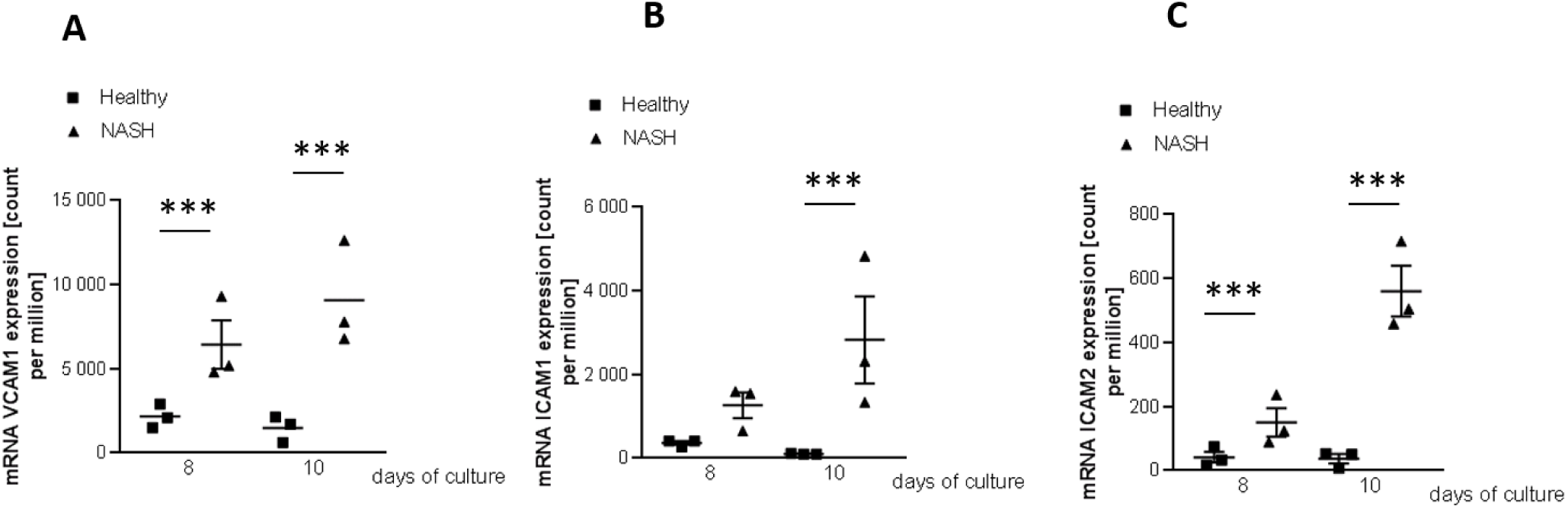
mRNA expression level. expressed in count per million at Days 8 and 10 in healthy and NASH condition with a scatter plot representation for VCAM1 (A), ICAM1 (B) and ICAM2 (C). *p< 0,050** p<0,010 ***p<0,001, DESeq2 test.

In conclusion, global gene expression analysis showed similar results to biochemical analysis. Together with the steatosis pathways observed under healthy conditions and maintained in the NASH 3D co-culture, inflammation and early induction of fibrosis are induced in the human primary 3D liver NASH model.

## Discussion

In our present study, we have set up an *in vitro* 3D NASH model by co-culturing four human primary liver cell types including hepatocytes, stellate, endothelial and Kupffer cells. The major challenge was to develop a medium able to maintain these four primary cell types in culture, during an extended period of time and to preserve PHH mature phenotype.

Our custom Liver Co-Culture media, optimized for 3D co-culture, enabled us to successfully ensure proper cell viability up to 2 weeks *in vitro*. Transcriptomic analyses performed at Day 10 indicated that PHH were still polarized and functional with the detection of ABCC2, ABCB11, CYP3A4, and CYP7A1 gene expression.

In NASH, injured steatotic hepatocytes induce an inflammatory environment leading to HSC activation and fibrosis. Since this mechanism results from hepatocyte ballooning, reflecting cell death and that the 3D NASH model requires a sustained viability to perform analysis, an inflammatory stimulus was provided by adding TNFα cytokine to the NASH culture media. PHH injury was mainly characterized by a decrease of CYP3A4 activity and mRNA expression at day 10 (Woolsey et al., 2015).

Intriguingly, the 3D liver co-culture exhibits high basal levels of triglycerides, as well as significant lipid droplet staining related to steatosis. An enrichment of pathways governing HSC activation and promoting early fibrosis is also observed.

Thus, the human 3D liver co-culture model displays steatosis and an activation of HSC leading to fibrosis, which are two main characteristics of NAFL/NASH. This model is able to respond to the inflammatory environment and to enhance the expression of proinflammatory cytokines like IL-6, as described in the literature (Bocsan et al., 2017, Rabelo et al., 2010). Several studies associated an increased CCL2 chemokine with steatohepatitis in chronic hepatic injury through an enhancement of pro-inflammatory monocyte/macrophage influx in the liver (Baeck et al., 2012, Tosello-Trampont et al., 2012, Narayanan et al., 2016, Krenkel and Tacke, 2017). Thus, CCL2 is thought to link steatosis and inflammation, and accordingly, its expression is upregulated in the human 3D NASH model.

An increase of CXCL8 and CXCL10 is also observed in the 3D liver NASH model confirming the induction of an inflammatory environment, which is a key characteristic of NASH disease (Koyama and Brenner, 2017). As previously described, CXCL8 could be a marker of NASH combined with diabetes (Estep et al., 2009). Recently, Zimmermann et al. established a positive correlation between CXCL8 mRNA expression and liver fibrosis stages showing that a high level of transcript is found in a mouse model of severe F4 fibrosis. Furthermore, an upregulation of CXCL8-binding CXCR1 and CXCR2 receptors has been positively correlated with chronic liver diseases (Zimmermann et al., 2011). CXCL10 cytokine expression is also correlated with fibrosis score (Domagalski et al., 2015). The pivotal role of CXCL10 in NASH was also shown *in vivo* using CXCL10 deficient mice, since a decrease in liver steatosis, injury, inflammation and fibrosis have been reported in this model compared to wild type animals (Zhang et al., 2014).

The increase of CXCL10 mRNA and secreted protein in 3D co-culture maintained in the NASH-like environment together with the key role of CXCL10 in NASH disease reported in the literature, underlying the relevance of our human 3D NASH model (Zhang et al., 2014, Tomita et al., 2016).

Fibrogenesis is known to be associated with the synthesis and the activity of matrix metalloproteases that regulate extracellular matrix (ECM) turnover during hepatic fibrosis. Among them, MMP2 and MMP9 are induced in fibrotic livers and are involved in the early disruption of the ECM in “pathologic” liver. Moreover, up-regulation of MMP2 was identified in human fibrotic liver, whereas MMP9 induction has been highlighted in the rodent NASH model (Okazaki et al., 2014, Giannandrea and Parks, 2014, Robert et al., 2016, Friedman, 2003). Later signals, such as collagen formation, are not observed probably resulting from a 10-day co-culture limitation.

The translatability is further strengthened by the differential regulation of the serum endothelial dysfunction markers ICAM-1 and VCAM-1. These two markers are of major interest to validate the induced fibrosis in the NASH model. Indeed, ICAM-1 was found to be significantly higher in serum from NASH patients compared to serum of NAFL and healthy patients (Ito et al., 2007), supporting the role of ICAM-1 as a potential disease progression biomarker. Regarding VCAM-1, it has also been recently validated as an accurate biomarker of fibrosis in NASH patients (Lefere et al., 2017). Furthermore, apoptosis signaling kinase 1 (ASK1/MAP3K5) has been largely described in the literature as a pharmacological target for NASH disease treatment (Povsic et al., 2019, Connolly et al., 2018, Drescher et al., 2019) and, interestingly, its up-regulation is observed in our human 3D NASH model (Table S2).

Finally, PD-L1 (CD274) gene expression has been shown to be enhanced in the human 3D NASH model and this finding is relevant with the induction of fibrosis. An up-regulation of PD-L1 gene has been described in injured liver and associated with HSC immunomodulatory activity (Yu et al., 2004) and hepatocyte damage leading to inflammatory processes (Wu et al., 2012). The induction of PD-L1 and PDCD1LG2 in the human 3D NASH model suggests that this pathway could qualify as an attractive avenue for NASH treatment.

In conclusion, we have developed a 3D NASH model with 4 human primary cell-types which include liver sinusoidal endothelial cells for the first time. These cells were shown to play a pivotal role in NAFLD/NASH progression from the simple steatosis to the early NASH stage probably by activating HSC and KC cells (Miyao et al., 2015, Ramachandran et al., 2019). Optimization of the model, for example by using microfluidic devices, could be an option to increase cell survival and to mimic chronic disease. However, it will likely not be suitable for screening purposes.

Our 3D NASH model could be maintained for 10 days *in vitro* and it showed triglyceride content leading to steatosis, an inflammatory response and activation of fibrosis-related pathways that are also associated with NASH. The relative extended life time of the 3D model culture makes it an attractive platform to evaluate preventive and curative treatments with drug candidates. These experiments will be reported in due course.

## Materials and Methods

Liver microenvironment under NAFLD and NASH diseases includes many circulating risk factors, which were incorporated into cell media to promote *in vitro* NAFLD or NASH-relevant phenotypes. Those risk factors include high glucose and insulin concentrations, excess of free fatty acids (FFAs) and endotoxins. For steatosis induction, FFAs (palmitic and oleic acids) were used which lead to the accumulation of intracellular lipid droplets. This lipotoxic phenotype is the most commonly used stimuli for NAFLD *in vitro* models, as it can also lead to an increase of inflammatory cytokines levels (Alkhouri et al., 2009, Miyao et al., 2015). TNFα was added to induce inflammation process and activate Kupffer cells. Cells were embedded in a hydrogel of rat collagen in 96 wells plate and NASH environment was induced with a media containing FFAs and TNFα. Considering the main characteristics of NASH pathology, hepatocytes injury, steatosis, inflammation and fibrosis were assessed biochemically and via transcriptomics.

### Cells 3D co-culture in hydrogels/matrix of collagen

Primary Human Hepatocytes (PHH) were obtained from LONZA (Walkersville, Maryland, USA) and Human Stellate Cells (HSC), Kupffer Cells (KC) and Liver Endothelial Cells (LEC) were provided by SAMSARA (San Diego, California, USA). PHH, HSC and LEC cells were seeded in red phenol free William E medium (Gibco ref. A12176) supplemented with 5% Fetal Bovine Serum, primary hepatocyte thawing and plating supplements solution (ThermoFisher scientific, ref. CM3000) and 1% of a Non-Essential Amino Acids solution (Gibco ref.11140050). PHH, LEC and HSC primary cells were embedded at 0.5 10^6^, 0.1 10^6^ and 0.1 10^6^ cells/mL, respectively in a half RAFT™3D collagen hydrogel (LONZA ref.016-0R92) in 96 wells plate as recommended by the provider. Cells were co-cultured in DMEM low glucose, pyruvate, no glutamine, no red phenol (Gibco ref.11054) supplemented with Bovine Serum free fatty acid free 0.125% (Sigma Aldrich, ref.A7030), 50U/mL penicillin/streptomycin (Gibco, ref.15140122), dexamethasone 0.1µM, ITS-G 1X (Gibco, ref.41400045), GlutaMAX™ 1X (Gibco, ref.35050061), HEPES 15mM (Gibco, ref.15630080), Non-Essential Amino Acids solution 1X (Gibco, ref.11140050), Acid L-ascorbic 2.5 mg/mL (Sigma, ref.A4403) and Glucagon 0.1µg/mL (Sigma, ref.G2044). This liver co-culture medium is named healthy media. After three days of culture, co-cultures were incubated either in healthy media or in a media mimicking NASH environment supplemented with glucose 25mM (Sigma), Oleate acid at 40µM (Sigma), Palmitate acid at 60µM (Cayman Chemical) and TNFα at 5 ng/mL (PeproTech). Healthy and NASH medium were changed every 2/3 days. At Day 6, Kupffer cells were added to the co-culture at 0.2 10^6^ cells/mL. Supernatants and embedded cells were sampled on days 3, 6, 8, 10, 13 and 15 for analysis.

### Viability and hepatocyte metabolism

Viability was assessed by ATP measurement using CellTiter-Glo 3D assay (Promega ref. G9681). PHH metabolism was measured through CYP3A4 activity with Luciferin-IPA CYP3A4-P450 Glo assay (Promega, ref. V9002).

### 3D primary liver cell co-culture immunostaining

3D hydrogels were fixed with 4% PFA/PBS during 15min at RT and immunostainings were performed. PHH cell membrane was stained with a rabbit anti-cytokeratin 18 antibody (EPR17347, Abcam #Ab181597) and an Alexa Fluor 594 conjugated goat anti-rabbit IgG secondary antibody (Molecular probe #A11012). HSC were stained with a mouse anti-αSMA antibody (Sigma #A5228) and an Alexa Fluor 680 conjugated goat anti-mouse IgG secondary antibody (Molecular probe #A21057). KC were stained with an FITC conjugated anti-CD68 antibody was used (KP1, Sigma #FCMAB205F), nuclei were labeled using NucBlue live (Thermo Fisher Scientific, #R37610). Image acquisitions were performed with a Leica SP8X confocal microscope, 40X water objective, Z stack of Z= 30µm zoom 0.8. 3D reconstitution was performed with IMARIS software 9.1.2.

### Triglyceride content

Triglycerides content was measured on co-culture supernatants with PicoProbe Triglyceride quantification assay (Biovision ref. K614-100). Intrahepatic lipid droplets (LD) were stained with lipidTox Green probe (Thermo molecular probe ref.H34475) and PHH cell membrane with CK18 immunostaining. Images of 3D co-cultures were acquired with a LEICA SP8X confocal microscope with a 40X water immersion objective and a z-stack of z= 30 µm, zoom 0.8. 3D reconstitution was performed with IMARIS software 9.1.2.

### Cytokine/metalloprotease release

Cytokine in co-culture supernatants were measured using MSD Technology with the U-plex biomarker human group 1 kit (MSD technology ref. K15067L-2). Samples were diluted at 1/4 for IL-6, CXCL8 and CCL2 measurement and no dilution was performed for CXCL10 quantification. Secreted metalloproteinases MMP2 was quantified with the human MMP2 ultra-sensitive kit (ref. K151FYC-2) from MSD Technology with a ½ dilution.

### Statistical methodology of biochemical parameters

To compare NASH and healthy models, a two-way ANOVA was performed on each parameter with group treatment (Healthy or NASH), day and their interaction as fixed-effects factors, and with experiment and interaction of experiment, group and day as random effect factor. Comparisons NASH versus healthy were provided for each day and no correction for multiplicity was done. To analyze the kinetic of the healthy model, a one-way ANOVA was performed on each parameter with day as fixed-effect factor and experiment and experiment by day as random effect factor. A Bonferroni-Holm’s correction was applied on p-values to compare: Day 6 vs Day 3; Day 8 vs Day 6; Day 10 vs Day 8 for comparison of both conditions and Day 10 vs Day 13 and Day 13 vs Day 15 for healthy time course assessment. Either a log or rank transformation was applied on studied parameters and fold changes were calculated. For log-transformed variables, the differences estimated from the model and their confidence intervals were back-transformed by using an exponential function. For rank-transformed variables, fold change and confidence intervals were estimated by Hodges-Lehmann’s method.

### RNA extractions

Hydrogels embedded co-cultures containing 0.2 10^6^ cells by well, were rinsed twice with cold PBS and stored at −80 °C until use. For RNA extraction, two wells were lysed in a final volume of 700 μl of Qiazol lysis reagent (Qiagen). Lysates were homogenized using CK14-2 ml tubes in Precellys tissue homogenizer instrument (Bertin). The lysats were collected and 140 μl of chloroform were added, vortexed and centrifuged at 12000 rpm for 15 minutes at 4 °C. Recovered aqueous phase was processed with an on-column DNAse treatment and a RNeasy mini kit (Qiagen) as recommended by the provider. RNAs were recovered in 30 μL of RNase free water. RNA quality was controlled on RNA LabChip with a 2100 Bioanalyzer (Agilent). Samples with RIN>7.4 were further processed and their concentration was quantified by Xpose spectrophotometer (Trinean).

### RNA libraries and sequencing

RNA-Seq libraries were generated with 15 ng of total RNA. cDNAs were generated with SuperScript VILO cDNA Synthesis Kit (Thermo Fisher) and the following steps for library construction were performed using the “AmpliSeq for Illumina Transcriptome Human Gene Expression Panel” according reference guide (Illumina).

Libraries were quantified and qualify respectively by Qubit (Invitrogen) and Bioanalyser (Agilent. Libraries were pooled an equimolar concentration at 4 nM, denatured and diluted at a final concentration of 1.4 pM. Sequencing was performed on Illumina NextSeq500 with NextSeq 500/550 High Output v2 kit and sequencing parameters of 2 × 151 bp Pair end, dual index (2 × 8bp).

Generated raw files were converted into FASTQ files and were analyzed on Array studio (V10.0.1.81 (Omicsoft, Qiagen).

Briefly, raw data QC was performed then a filtering step was applied to remove reads corresponding to rRNAs as well as reads having low quality score. Mapping and quantification were performed using OSA4 [1C] (Omicsoft Sequence Aligner, version 4)(Hu et al., 2012). Reference Human genome Human.B38 was used for mapping and genes were quantified based on RefSeq gene annotations. Differentially expressed transcripts were identified with DESeq2 (Love et al., 2014). The variable multiplicity being taken into account and false discovery rate (FDR) adjusted p-values calculated with the Benjamini-Hochberg (BH) correction (Benjamini and Hochberg, 1995). Significant DEGs were defined as p < 0.05 after adjustment for false discovery and average fold change between condition replicates of >1.8. DEGs were further analyzed using Ingenuity Pathway Analysis (IPA) (QIAGEN Inc., https://www.qiagenbioinformatics.com/products/ingenuity-pathway-analysis) (Kramer et al., 2014).

All authors had access to the all data and have reviewed and approved the final manuscript.

## Supplementary Materials

**Fig. S1:**
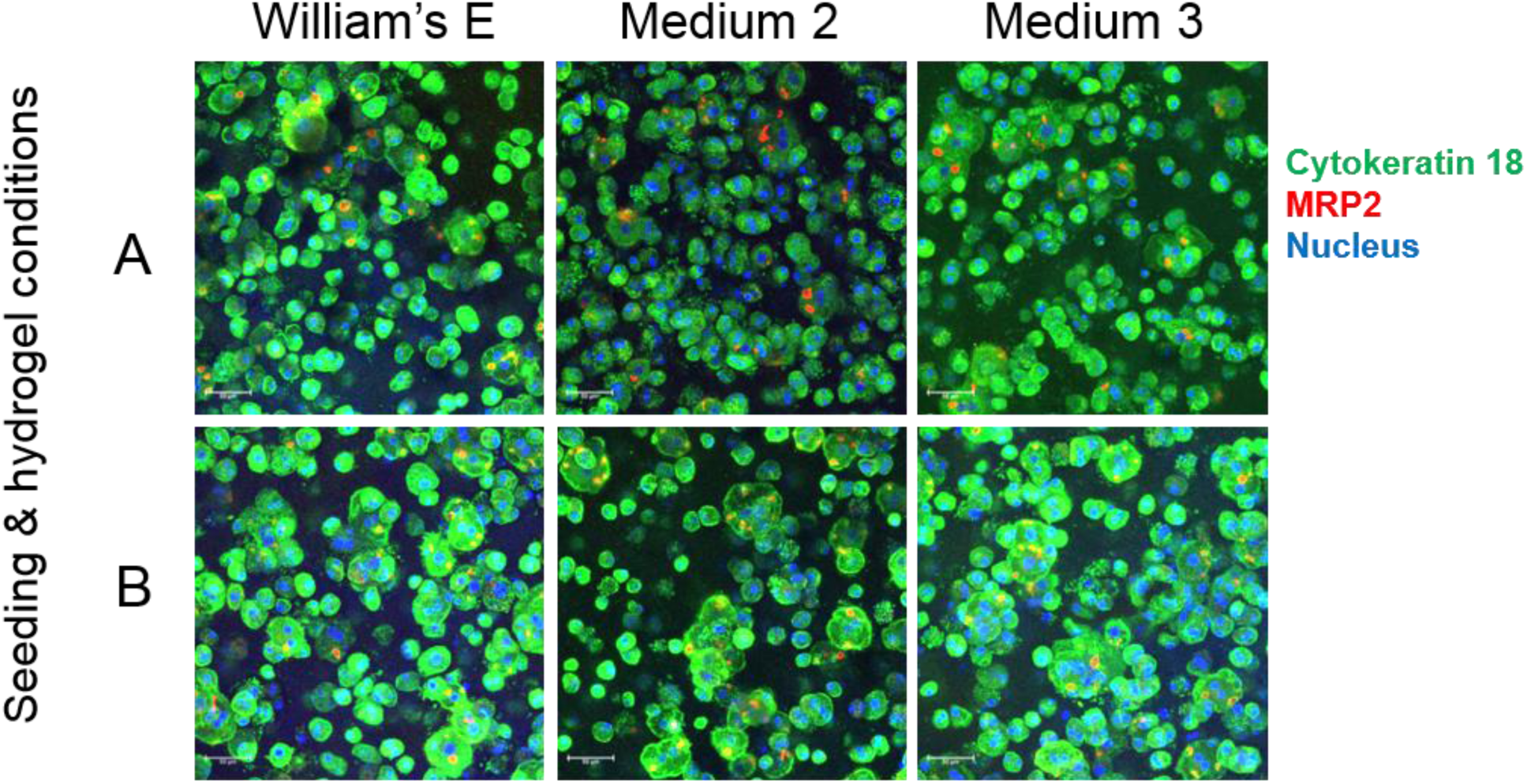
Seeding & hydrogel condition A & B: William’E media, medium 2 (Advanced DMEM) and medium 3 (DMEM low glucose) maintain PHH polarisation in culture for 14 days. Leica SP8X confocal microscope, 40X water objective, Z stack of Z= 30µm zoom 0.8 PHH staining of MRP2 in red, nuclear in blue and lipid droplet in green

**Fig. S2.**
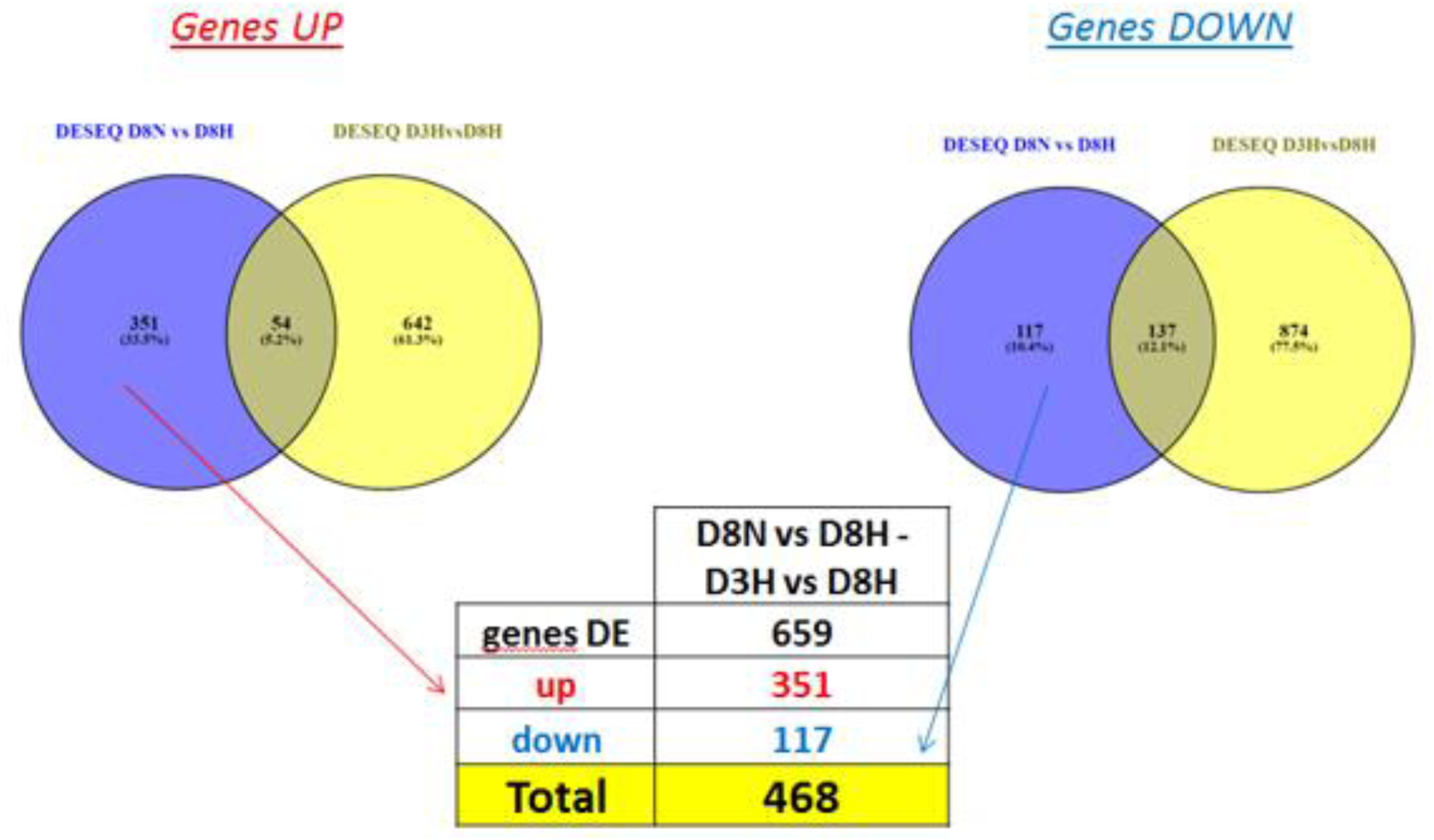
Differential expression analysis (DESeq): NASH vs Healthy at Day8 minus effect D3H vs D8H (separate effect up and down regulation) Cut off: abs Fold change > 1.8, FDR_BH <0.05

**Table S1.**
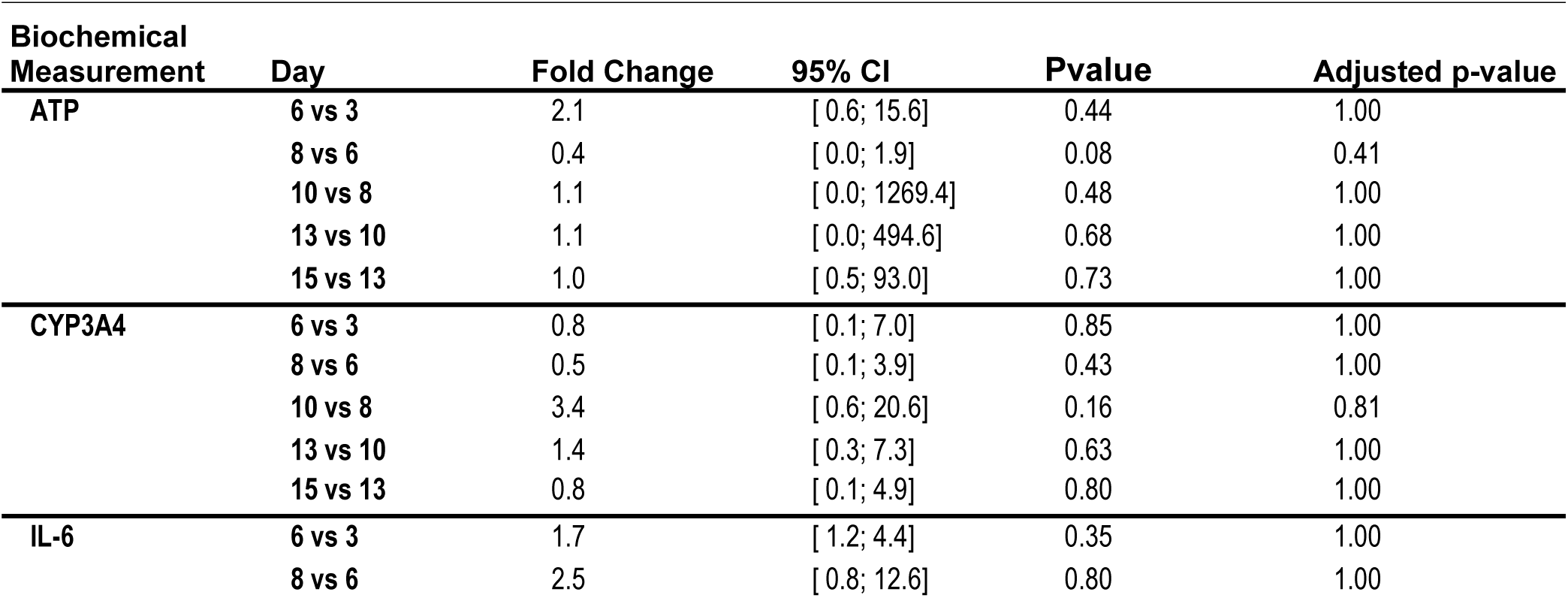
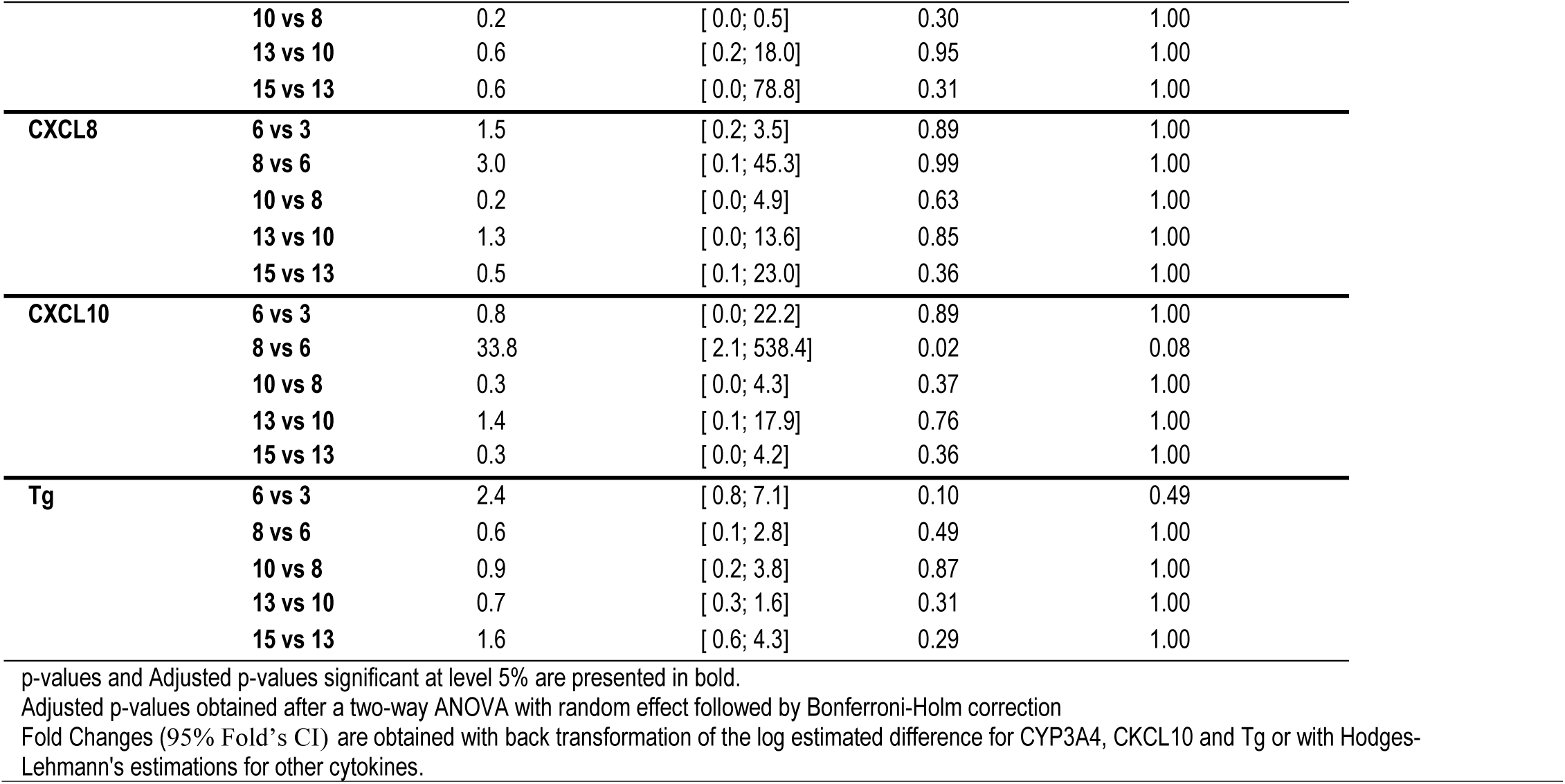
Pairwise comparisons of ATP, CYP3A4, cytokines and triglyceride content in 3D human liver co-culture

**Table S2.** Stability of 3D liver co-culture model: gene expression fold changes and p-values.

| Gene Name | Day 8 vs Day 3-fold change | Day 8 vs Day 3 p-value | Day 10 vs Day 3-fold change | Day 10 vs Day 3 p-value | Day 10 vs Day 8-fold change | Day 10 vs Day 8 p-value |
| --- | --- | --- | --- | --- | --- | --- |
| ABCB1 | 1.6 | 0.42 | 1.5 | 0.30 | 1.0 | 0.94 |
| ABCB11 | -1.4 | 0.54 | -1.2 | 0.71 | -1.2 | 0.29 |
| ABCC2 | 1.4 | 0.58 | 1.5 | 0.29 | -1.1 | 0.54 |
| ACTA2 | 2.8 | **3.68E-14** | 2.4 | **4.739E-12** | 1.2 | 0.16 |
| COL1A1 | 1.2 | 0.44 | 1.4 | 0.06 | -1.2 | 0.24 |
| CXCL10 | 5.3 | **1.03E-06** | 6.0 | **1.615E-10** | -1.1 | 0.36 |
| CXCL8 | -3.4 | **2.9E-10** | -3.2 | **4.894E-12** | -1.1 | 0.70 |
| CYP3A4 | -6.2 | **2.79E-08** | -4.7 | **1.895E-08** | -1.3 | **0.04** |
| CYP7A1 | -1.6 | 0.36 | -1.1 | 0.85 | -1.4 | **0.004** |
| IL-6 | -1.1 | 0.87 | -1.1 | 0.83 | -1.0 | 0.97 |
| UGT1A1 | -2.1 | 0.10 | -1.9 | 0.06 | -1.1 | 0.59 |

Fold change of count per million values and p-value calculated with DESeq2. Significant Differential Expression Gene (DEG) are defined as p<0.05 in bold.

**Table S3.**
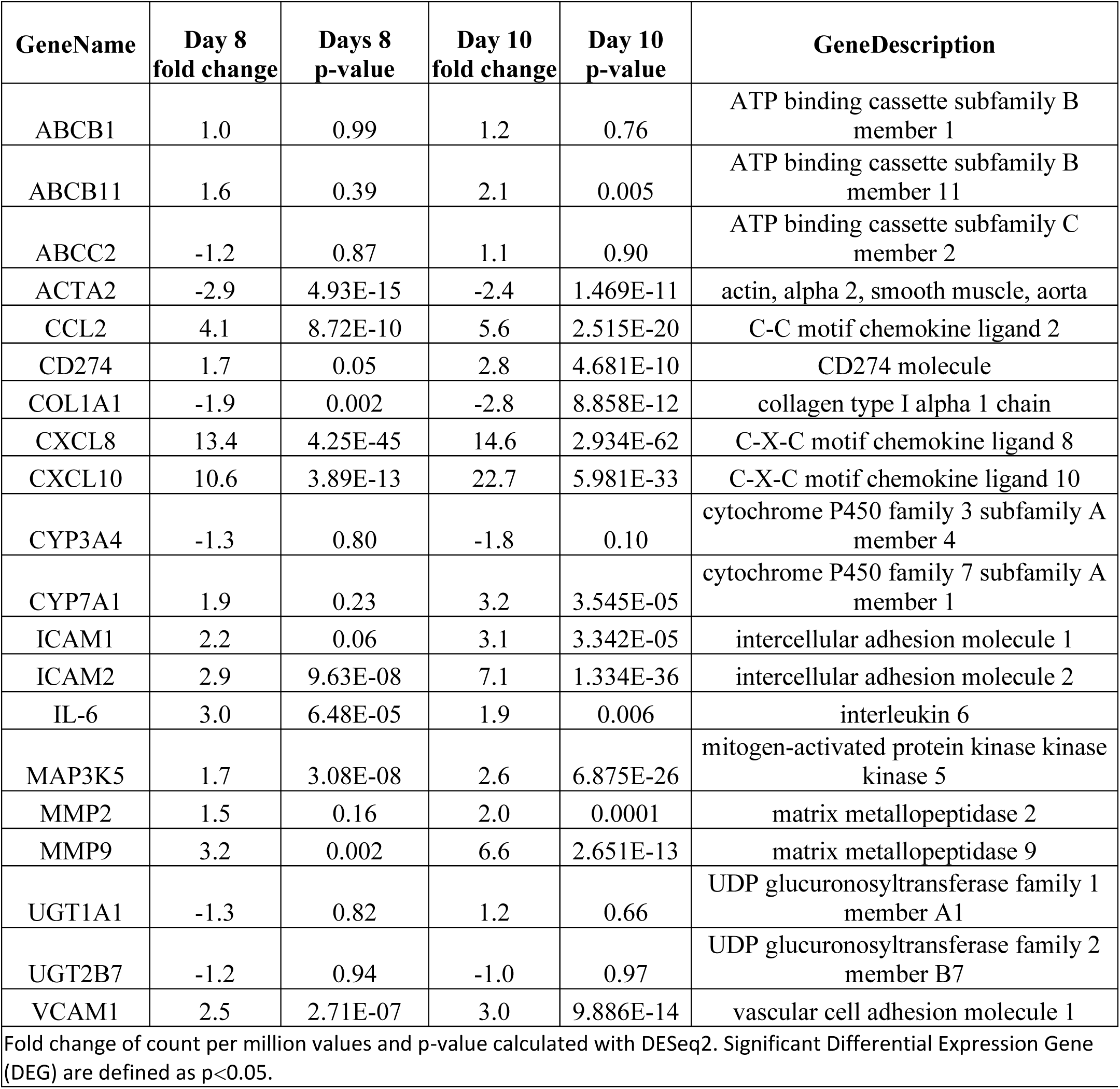
Fold change and p-value of mRNA gene expression in NASH as compared to Healthy conditions at Day 8 and Day 10

**Table S4.** – Fold change IL-6, CXCL8, CXCL10, CCL2 and MMP2 secretions between healthy and NASH 3D co-culture at Day 6, Day 8 and Day 10

| Cytokine | Day | Fold<br>Change | 95% CI | Pvalue |
| --- | --- | --- | --- | --- |
| IL-6 | 6 | 5.0 | [ 2.2; 7.4] | 0.01 |

| **Cytokine** | **Day** | **Fold Change** | **95% CI** | **Pvalue** |
| --- | --- | --- | --- | --- |
|  | 8 | 1.3 | [ 0.0; 6.0] | 0.23 |
|  | 10 | 1.7 | [ 0.0; 5.0] | 0.47 |
| **CXCL8** | 6 | 10.3 | [ 1.0; 129.0] | **0.02** |
|  | 8 | 4.7 | [ 0.0; 117.6] | **0.02** |
|  | 10 | 9.6 | [ 0.0; 33.5] | 0.09 |
| **CXCL10** | 6 | 32.5 | [ 1.3; 817.9] | **0.04** |
|  | 8 | 8.4 | [ 0.4; 157.6] | 0.14 |
|  | 10 | 22.6 | [ 1.2; 425.3] | **0.04** |
| **CCL2** | 6 | 8.7 | [ 0.8; 98.6] | **0.03** |
|  | 8 | 3.8 | [ 0.0; 6401.5] | 0.19 |
|  | 10 | 3.4 | [ 0.0; 194.8] | 0.30 |
| **MMP2** | 6 | 1.2 | [ 0.0; 192.8] | **0.02** |
|  | 8 | 1.7 | [ 0.3; 6.9] | **0.0005** |
|  | 10 | 1.4 | [ 0.3; 6.7] | **0.0003** |

p-values significant at level 5% are presented in bold. p-values obtained after one-way ANOVA with random effect followed by comparisons at fixed day. Fold Changes (95% Fold's CI) are obtained with back transformation of the log estimated difference for CXCL10 and with Hodges-Lehmann's estimations for other cytokines

## Acknowledgments

The authors want to thank Prof. François Pattou and Prof. Bart Staels for support and advice, and Federation Hospital & University INTEGRA who in association with SANOFI are part of the PreciNASH consortium.

## Author contributions

Agnes Jacquet and Lucile Hoet were responsible for the study design, data acquisition and biochemical data analysis; Sandrine Roche, Marie-Dominique Bock and Corinne Rocher were responsible for transcriptional data acquisition and analysis; Gilles Haussy was responsible for imaging data; Xavier Vigé and Zsolt Bocskei were responsible for critical revision of the manuscript; Tamara Slavnic and Valérie Martin were responsible for statistical analysis; Franck Augé, Jean-Claude Guillemot, Michel Didier, Aimo Kannt, Cécile Orsini and Vincent Mikol were responsible for critical revision of the manuscript and funding acquisition; Marion Duriez and Anne-Céline Le Fèvre were responsible for study supervision, study design, data acquisition analysis and interpretation and drafting the manuscript.

